# USP-ddG: A Unified Structural Paradigm with Data Efficacy and Mixture-of-Experts for Predicting Mutational Effects on Protein-Protein Interactions

**DOI:** 10.1101/2025.11.09.687124

**Authors:** Guanglei Yu, Xuehua Bi, Qichang Zhao, Jianxin Wang

## Abstract

Accurately estimating changes in binding free energy (*ΔΔG*) is essential for understanding protein-protein interactions (PPIs) and guiding rational protein design. Existing deep learning approaches benefit from pre-training on large structural datasets, but they often require heavy computation and overlook data efficacy. In addition, prior models typically assume fixed backbone structures and ignore conformational dynamics, limiting generalization. We present USP-ddG, a unified structural paradigm for *ΔΔG* prediction. USP-ddG integrates a dual-channel architecture with three complementary components: (i) inverse folding-based log-odds ratio, (ii) empirical energy terms from FoldX, and (iii) a geometric encoder with Gaussian noise to capture relaxed conformations. To enhance representation power, we introduce a framework that integrates feed-forward network (FFN) and Mixture-of-Experts (MoE) to model domain-invariant and -specific features, respectively. We further propose CATH-guided Folding Ordering (CFO), a data efficacy strategy that organizes samples to mitigate catastrophic forgetting and data distribution bias. USP-ddG consistently outperforms existing state-of-the-art (SoTA) methods on the SKEMPI v2.0 benchmark, including the challenging hold-out CATH test set. It achieves superior accuracy on both single- and multi-point mutations and demonstrates strong performance in antibody affinity optimization against H1N1 and HER2, and in assessing SARS-CoV-2 variants to hACE2. Ablation studies confirm the benefit of each component. These results highlight USP-ddG as a robust and data-efficient framework for modeling mutational effects on PPIs.

**Availability:** USP-ddG is publicly available at https://github.com/ak422/USP-ddG.

## 1 Introduction

As the ultimate executors of cellular structures and functions, proteins communicate through specific physical interactions and assemble into molecular machines, participating in nearly all cellular activities [13,21]. Mechanistic dissection of PPIs has provided a foundation for engineering therapeutics that aim at treating, for example, amyloid-related diseases [12,15] and cancers [22,33].

The binding between protein molecules can be regarded as a reversible and rapid equilibrium process, with the binding strength typically quantified by thermodynamic parameters, such as equilibrium dissociation constant (*K*_*D*_) or binding free energy (*ΔG*). Thus, developing methods to predict the changes in binding free energy (*ΔΔG*) caused by amino acid mutations is of critical importance in protein design [27]. This involves screening mutant sequences to identify optimized variants that significantly reduce *ΔG* and thereby enhance binding strength. However, the potential sequence space of mutations is extremely large, and conventional wet-lab experiments face significant challenges due to limited throughput and high costs. In this context, the development and application of computational methods capable of accurately and efficiently predicting *ΔΔG* have become an urgent need to bridge this gap [28].

Currently, a variety of computational approaches have been developed for predicting *ΔΔG*. However, traditional methods based on physical energy functions and empirical force fields, as well as experimental techniques, often suffer from inherent limitations such as operational complexity and low throughput. Among the SoTA computational protocols, flex ddG in the Rosetta suite estimates *ΔΔG* through sampling conformational ensembles and incorporating the “backrub” algorithm to account for protein conformational plasticity [4]. Although physically rigorous, this method suffers from low computational efficiency, with screening efficiencies up to five orders of magnitude lower than modern deep learning based approaches [5,45]. In contrast, FoldX utilizes its empirical effective energy function (EEEF) to rapidly and quantitatively assess the effects of mutations (*ΔΔG*) [11,14,34]. The parameters of this energy function are optimized through fitting to large-scale experimental datasets using machine learning techniques, rendering it highly reliable for protein engineering applications. Recently, energy terms derived from FoldX have been integrated as effective prior knowledge into deep neural networks, enabling fast and accurate prediction of *ΔΔG* [45].

With the rapid advancement of deep learning, data-driven approaches have emerged as promising alternatives for *ΔΔG* prediction [35,42,45]. Despite continuous improvements, predictive performance remains constrained by the scarcity of high-quality experimental annotations. To address this limitation, self-supervised pre-training on large-scale unlabeled structural data has become a prevalent paradigm for enhancing model generalization. Common proxy tasks for pre-training include diverse learning objectives, such as inverse folding modeling [44], atomic coordinate reconstruction [25], side-chain conformation prediction [24,26], and masked language modeling [43]. These pre-training strategies enable the learning of sequence and structure representations imbued with physicochemical constraints, while simultaneously highlighting the critical challenge of effectively transferring such prior knowledge to downstream *ΔΔG* prediction task [19].

In response, and under the assumption of fixed backbone upon mutations, BA-DDG estimates *ΔΔG* using the Boltzmann distribution, incorporating the difference in sequence negative log-likelihood scores derived from inverse folding model ProteinMPNN [10], thereby achieving substantially improved predictive accuracy [19]. In contrast, EBM-DDG relaxes the fixed backbone assumption to accommodate structural rearrangements upon mutations. It decomposes *ΔΔG* into a sequence term, derived from BA-DDG, and a structural term. The structural term is estimated using the diffusion-based score-matching model DSM-Bind [20], which generates mutant conformations and quantifies the associated energy changes, enabling more comprehensive *ΔΔG* modeling [38]. However, it overlooks the intrinsic conformational dynamics of proteins, and focuses on backbone-only determinants of *ΔΔG*, which may result in the loss of critical information contributed by conformational dynamics and side-chains.

In this work, we propose USP-ddG, a unified structural paradigm for predicting *ΔΔG* on PPIs. The model integrates both protein sequence and structure as input and decomposes *ΔΔG* into three components along with an auxiliary task. The overall architecture of our approach is structured as follows:

1. Component 1: leveraging the thermodynamic cycle definition, we use the inverse folding model Protein-MPNN to compute sequence negative log-likelihood scores, enabling USP-ddG to learn the correlations between residue preferences and mutational effects.
2. Component 2: this component focuses on modeling *ΔΔG* prediction with energy terms generated by force field FoldX, which enables the model to learn physical inductive bias from empirical data, providing to account for the thermodynamic properties of proteins.
3. Component 3: we employ an improved CATH-ddG of our previous work for encoding the protein sequence and 3D structure. To simulate structural dynamics, Gaussian noise is added to atomic coordinates during training. Specifically, for the improved CATH-ddG, we introduce a hybrid architecture that integrates FFN and MoE to capture domain-invariant and domain-specific features, respectively. In addition, we propose CATH-guided Folding Ordering (CFO), a curriculum learning-based strategy to enhance learning efficiency by optimizing the ordering of training data. Unlike random shuffling or order-fixed sampling, CFO aims at mitigating issues such as catastrophic forgetting and data distribution bias.
4. Integration of CATH domain alignment: to integrate protein domain classification information for better representation learning, we formulate the structural constraints derived from CATH domains as an auxiliary multi-label classification task within a self-supervised learning (SSL) framework, enabling the model to learn the structural clusters and functional families within its CATH superfamilies.

We conduct extensive experiments to validate the effectiveness of USP-ddG. Specifically, we adopt the independent hold-out CATH test set introduced in CATH-ddG [45], which is curated from SKEMPI v2.0 [18]. This independent test set contains 813 automatically constructed mutations, which ensures no overlap in CATH superfamily with the training set while also maintaining a maximum TM-score *<* 0.6 for structural similarity. Our results demonstrate that USP-ddG consistently achieves the best performance. In addition, ablation studies confirm the critical contributions of data efficacy and MoE strategies, as well as the use of inverse folding and improved CATH-ddG in enhancing model performance. Overall, the results strongly support the robustness and superior effectiveness of USP-ddG in predicting *ΔΔG* in a unified structural paradigm. Independent case studies demonstrate successful enhancement of binding affinity on antibody variants of CR6261 against H1N1 subtype of human influenza A pandemic, variants of trastuzumab against human epidermal growth factor receptor 2 (HER2), and variants in receptor-binding domain (RBD) of SARS-CoV-2 to human angiotensin-converting enzyme 2 (hACE2) receptor.

## 2 Method

### 2.1 Notation and Problem Definition

A protein chain 𝒜 is represented by *n* amino acids 𝒮_𝒜_ ∈ {1, …, 20 }^*n*^, and its atomic coordinates 𝒳 _𝒜_ ∈ ℝ^*n×m×*3^. Here, *m* represents the number of atoms per amino acid. A protein complex 𝒜ℬ is formed by the binding of two chains, 𝒜 and ℬ, and exists in a dynamic equilibrium between its bound state 𝒜ℬ_bnd_ and unbound state 𝒜ℬ_unbnd_. We refer to the unmutated protein chain as the wild-type (wt) and the mutated chain as the mutant (mut). For simplicity, this section focuses on two-chain complexes, using the notation {𝒮_𝒜ℬ_, 𝒳 } to represent the sequence and its corresponding structure.

The binding free energy (*ΔG*) of a protein complex is defined as the difference in Gibbs free energy between the bound state (*G*_bnd_) and the unbound state (*G*_unbnd_): *ΔG* = *G*_bnd_ − *G*_unbnd_. Based on this definition, the change in binding free energy (*ΔΔG*) upon amino acid mutation is given by:

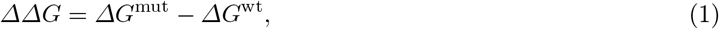

where *ΔG*^mut^ and *ΔG*^wt^ represent the binding free energies of the mutant 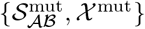 and wild-type 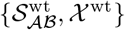 complexes, respectively. Our objective is to predict *ΔΔG* for a given set of mutations {wt → mut}.

### 2.2 Overview

In Fig. 1 (a), we present an overview of USP-ddG. The workflow consists of the following key steps: (i) Channel 1 (structure-fixed view; Equation 6) decomposes *ΔΔG* into an inverse folding component, log *p*( 𝒮_𝒜ℬ_ | 𝒳^FX^), derived from the negative log-likelihood scores output by ProteinMPNN, and an energy component, log *p*( 𝒳^FX^), calculated using the energy terms from FoldX. The prior work for estimating *ΔΔG* using the inverse folding model, as well as the derivation under the structure-fixed assumption, are provided in Appendices A and B, respectively. (ii) Channel 2 (structure-relaxed view; Equation 7) integrates our previous work, CATH-ddG, with an auxiliary SSL task that aligns the latent embeddings to protein domain induction preferences, thereby enhancing both predictive performance and interpretability in downstream *ΔΔG* prediction. Finally, the loss function is defined as the weighted sum of above supervised learning and SSL tasks by the learnable scalar parameters (Equation 11).

**Fig. 1:**
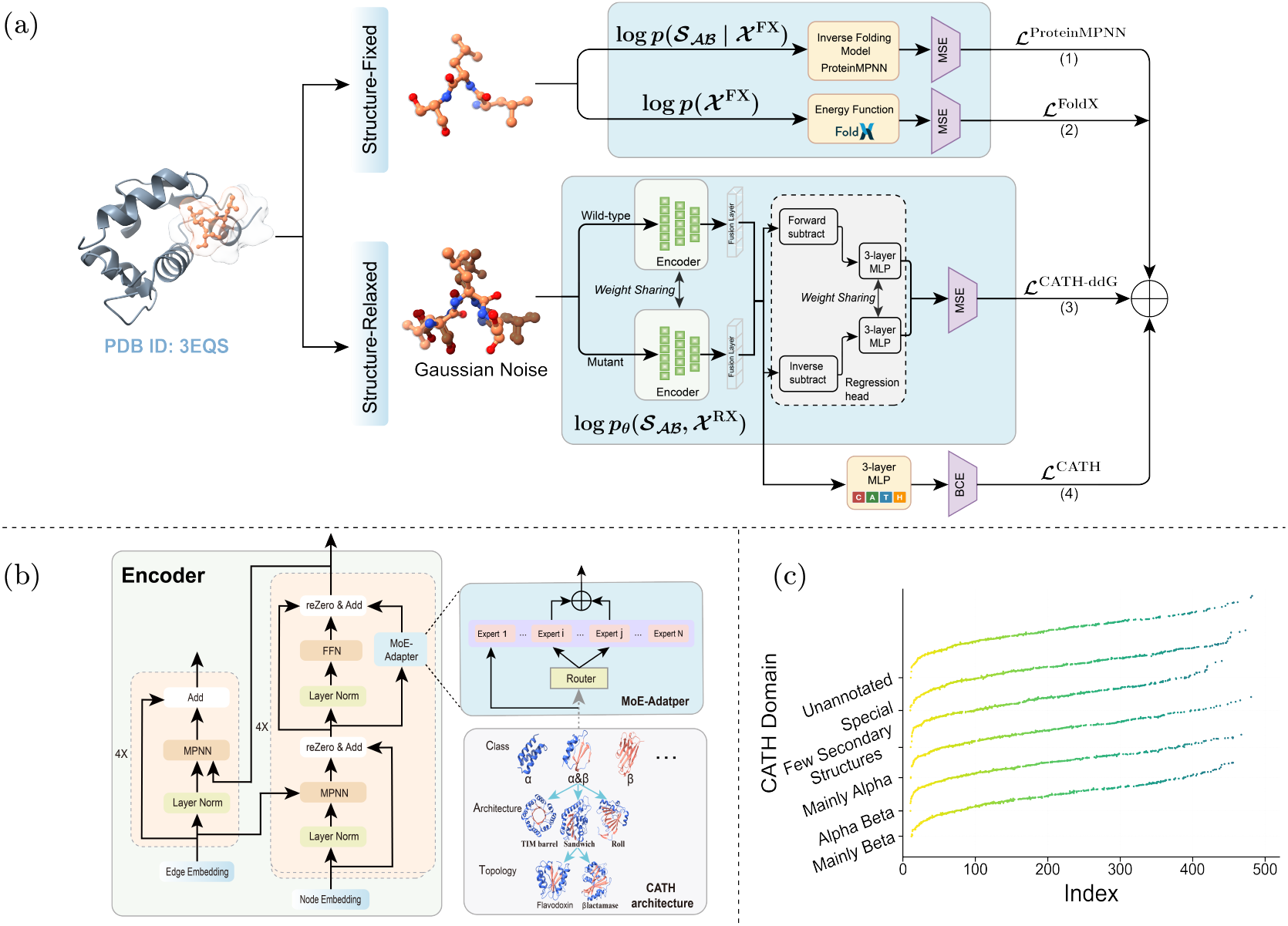
(a) Overview of the proposed USP-ddG framework. The framework is composed of a dual-channel and four task branches: (1) inverse folding-based supervised learning task, (2) energy function-based supervised learning task, (3) self-supervised learning task for CATH domains, and (4) training with structure noise-based supervised learning task. (b) To utilize the advantage of sparsely activated of MoE to capture the shared and distinctive features across diverse protein domains, we improved the Feed-Forward Network via FFN with MoE to learn domain-invariant and specific features of protein domains. (c) Illustration of CATH-guided Folding Ordering. These results are based on simulated data.

### 2.3 Modeling with Structure-relaxed Assumptions

Inverse folding-based models for predicting *ΔΔG* (e.g., BA-DDG and EBM-DDG) typically rely on two assumptions: (1) that protein function is primarily dictated by backbone atoms, and (2) that the protein structure exists in a unique, static conformation. However, protein function is orchestrated by a dynamic conformational landscape that integrates the contributions of both the rigid backbone (BB) and the flexible side-chains (SC). Consequently, prior approaches that rely exclusively on the protein backbone fail to capture the full physicochemical complexity of the system. To this end, we propose a unified *ΔΔG* prediction framework that incorporates side-chain atoms and dynamic conformations into the structural assumptions. To formalize this, we correspondingly define *ΔG* as:

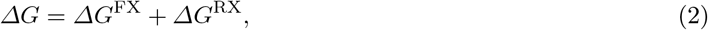

where *ΔG*^FX^ and *ΔG*^RX^ correspond to the binding free energy contributions under the structure-fixed (FX, unique conformation) and structure-relaxed (RX, dynamic conformation) hypotheses, respectively. Accordingly, we define the protein structure under each hypothesis as follows:

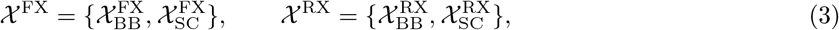

where 𝒳 ^FX^ denotes the protein structure under the structure-fixed hypothesis, defined by a unique backbone 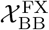 and side-chains 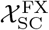. In contrast, 𝒳 ^RX^ represents the structure under the structure-relaxed hypothesis, characterized by a multi-conformation backbone 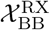 and side-chains 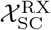.

#### Structure-fixed Assumption (*ΔG*^FX^)

Under the structure-fixed hypothesis, the binding free energy *ΔG*^FX^ can be derived from the Boltzmann distribution [2] and is expressed in terms of the probabilities of the bound state and the unbound state, denoted as 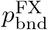 and 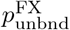, respectively, as shown below:

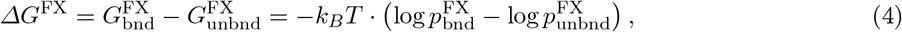

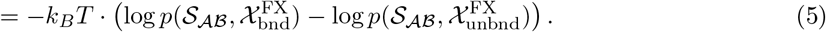

Applying Bayes’ theorem, similar to the EBM-DDG approach [38], this simplifies to:

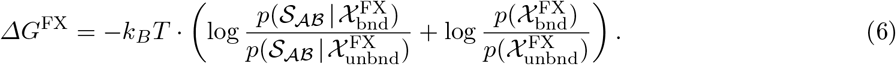

The probability of unbound state is approximated as 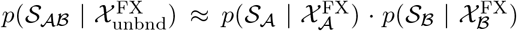. Furthermore, we leverage the common assumption in protein sequence design that the sequence is largely determined by the backbone structure. Thus, the conditional probabilities 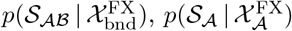, and 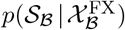 are provided by the pre-trained inverse folding model ProteinMPNN, denoted as *ΔG*_ProteinMPNN._ And the second term in Equation 6, 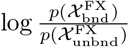, which corresponds to the binding free energy of the full complex, is estimated by EBM-DDG under the structure-fixed and backbone-only assumption using the diffusion-based score-matching model DSMBind [20]. In contrast, our method incorporates both backbone and side-chains, and approximates this energy term with the empirical force field FoldX, yielding *ΔG*_FoldX_.

#### Structure-relaxed Assumption (*ΔG*^RX^)

Following prior studies and assuming that the unbound state structure is well represented by the bound state conformation, the binding free energy under the relaxed structure hypothesis is formulated by a neural network model:

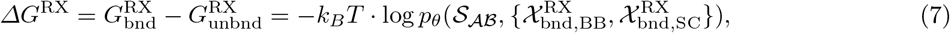

where 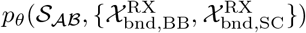 is parameterized by the deep learning neural network CATH-ddG, denoted as *ΔG*_CATH-ddG_, with learnable parameters *θ*, taking the *S*_*AB*_ and structure 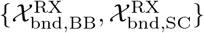 with Gaussian noise to computationally simulate its relaxed conformational state as input.

### 2.4 Unified Structural Paradigm

Under the preceding assumption, we employ a unified structural paradigm *ΔG* = *ΔG*^FX^ + *ΔG*^RX^ to achieve the evaluation of *ΔΔG*. This approach, named USP-ddG, integrates the derived components, thereby yielding the overall binding free energy *ΔG* as follows:

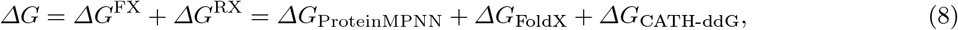

where the first term is predicted by the pre-trained inverse folding model ProteinMPNN, the second term is calculated using FoldX, and the third term is predicted by the improved, trainable neural network model CATH-ddG. Finally, the change in binding free energy *ΔΔG* is expressed as:

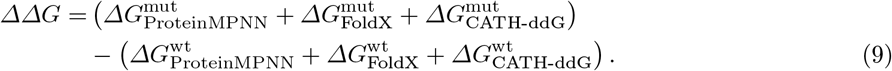

We follow the setup of Yu et al. [45] by incorporating an auxiliary self-supervised Binary Cross-Entropy (BCE) loss to align model training with the inductive biases of protein domains. In our experiments, −*k*_*B*_*T* is treated as a learnable parameter. The final objective function is characterized by the following composite Mean Squared Error (MSE) loss and Binary Cross-Entropy (BCE) loss:

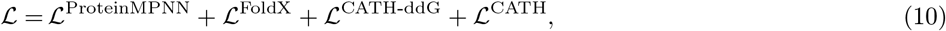

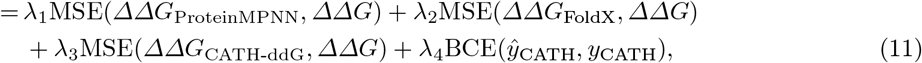

where 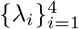 are learnable scalar parameters, as shown in Fig. 1 (a).

### 2.5 Training with Mixture-of-Experts

We leverage sparse MoE [17,37] to establish a scalable framework aimed at mitigating the issue of structural overlaps across different CATH superfamilies [7,30]. This section first outlines the MoE mechanism and then details its integration into the FFN layer.

The MoE contains two components: the *N* experts 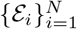 and a routing network ℛ. While it maintains efficiency by activating only a subset of experts for each input token, its output is combined dynamically via a weighted strategy by the router . Inspired by [46], we use the LoRA [16] adapter as the expert in MoE to speed up the adaptation on downstream tasks. Our MoE-Adapters are implemented in all the structured Transformer blocks of the spatial and sequential encoders of CATH-ddG, as shown in Fig. 1 (b). Formally, let 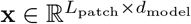 be the input tokens. The combined output 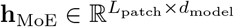 is then computed as:

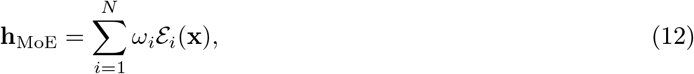

where 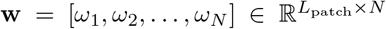 represents the gating weights assigned by router ℛ, dictating each expert’s contribution to the output **h**_MoE_. The gating weights are computed using a learnable matrix 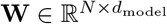 and a learnable matrix for the noise component 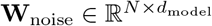 as follows [37]:

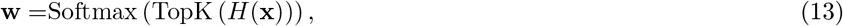

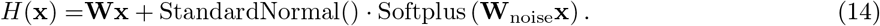

For the FFN layer, we adopt an architecture inspired by DeepSeek-MoE [8,23], which utilizes a hybrid layer of shared and routed experts. Specifically, the shared expert is dedicated to learning domain-invariant knowledge and reducing redundancy among the routed experts, while the routed experts specialize in domain-specific knowledge of protein structures. Accordingly, the output of FFN layer after the attention head, denoted as 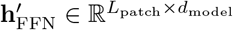, is subsequently redefined as:

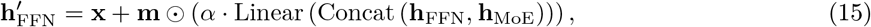

where **m** ∼ Bernoulli(*p* = 0.1) is the dropout mask vector, ⊙ is the Hadamard product operation, *α* is a learnable scalar for reZero [3], and Concat(·) represents the concatenation operation, as shown in Fig. 1 (b).

### 2.6 Training with Data Efficacy

Machine learning models typically achieve reliable performance when the training data accurately capture the real-world distributions. However, even SoTA algorithms often suffer from limited generalization performance when deployed in distribution shifts [39,40]. This raises a critical question: how can we systematically enhance model performance without altering the data scale or model architecture? The answer may lie not in the data itself, but rather in how it is utilized. Currently, data efficiency aims at curating data in terms of both quantity and quality to improve model performance. In contrast, data efficacy is concerned with maximizing performance by optimizing the organization and utilization of training data [9].

Additionally, investigators have demonstrated that while easy negative samples promote shortcut learning, hard negatives are essential for compelling models to learn complex features, thereby ultimately facilitating the discovery of biological mechanistic principles [32,40]. To build models that generalize across diverse real-world scenarios, it is crucial to challenge them with appropriately difficult data. To this end, we propose the CATH-guided Folding Ordering (CFO) strategy, which organizes training data by leveraging evolutionary insights from CATH protein domain annotations to present increasingly difficult samples. More specifically, to construct the training set enriched with hard negatives, we partition the training data into Class-disjoint subsets according to the Class-level categorical definition in the CATH database. The permutation function *π*_CFO_, which defines the training order, is formalized as follows:

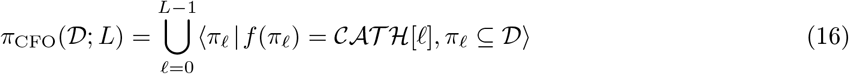

where *f* (*π*_*ℓ*_) returns the Class of *π*_*ℓ*_, 𝒞 𝒜𝒯ℋ = {0, 1, 2, 3, 4, 6} denotes the set of Class-level categories, 0 indicates unannotated entries [45], and *L* = |𝒞 𝒜𝒯ℋ |. Thus, based on the Class distributions, an illustration of CFO is provided in Fig. 1 (c), which starts with easier samples and progressively tackles harder ones.

## 3 Experimental Settings

This section introduces the experimental settings, including the datasets and baseline methods used for comparison. Comprehensive details regarding the evaluation metrics and parameter settings are provided in Appendices C and D, respectively.

### 3.1 Datasets

To train and evaluate USP-ddG, we utilize the largest available curated dataset, SKEMPI v2.0, which comprises 7,085 mutations. Previous studies have primarily benchmarked models using a PPI-disjoint cross-validation split on this dataset [19,24,26,38]. However, this evaluation strategy is susceptible to significant data leakage and may overestimate model performance. As demonstrated in our prior work [45], an average of 88.70% of mutation entries under this split are classified as easy mutations—defined as those exhibiting a maximum TM-score ≥ 0.6 when aligned to structures in the training set. To enable a more rigorous and functionally relevant assessment, we adopt a partitioning strategy based on CATH homologous superfamilies defined in CATH-ddG [45]. Thus, SKEMPI v2.0 is divided such that the hold-out CATH test set contains only complexes that share no CATH superfamilies with the training set. This procedure yields a challenging test set of 813 mutations, in which none are classified as easy mutations, thereby providing a robust evaluation of the model’s generalization towards unseen CATH homologous superfamilies.

### 3.2 Baselines

For the hold-out CATH test set selected from SKEMPI v2.0, we evaluate our two methods, USP-ddG^Random^ and USP-ddG, against three categories of SoTA baselines, utilizing retrained and retested results whenever available. Here, USP-ddG^Random^ and USP-ddG denote our approach trained on the randomly shuffled training data and the CFO training data 𝒟, respectively. Specifically, the first category is traditional energy function approaches, including flex ddG [4] and FoldX [11]. The second category is pre-training based approaches, including RDE-Network [26] and DiffAffinity [24]. The third category is supervised learning approaches, including CATH-ddG [45] and BA-DDG [19].

## 4 Results

We evaluated our algorithm using both random shuffling and the proposed CFO-based data efficacy strategy, benchmarking it against previous works. The results demonstrate that USP-ddG consistently and substantially outperforms USP-ddG^Random^ across all experiments. This consistent advantage confirms that the CFO-based strategy for organizing training data is significantly more effective than random shuffling. Detailed results are presented in the following sections. In the following tables, the best and second-best results are highlighted in **bold** and <u>underlined</u>, respectively. We also report the performance of USP-ddG for the binding of SARS-CoV-2 RBD to hACE2 in Appendix E.

### 4.1 Performance Comparison on SKEMPI v2.0 dataset

We evaluate our USP-ddG deep learning framework on the hold-out CATH test set curated from SKEMPI v2.0, and present the results in Table 1 for three categories of methods across seven evaluation metrics. To ensure robust assessment, five independent experiments with different random seeds are conducted. The key findings are summarized below: (1) USP-ddG consistently outperforms all baselines on both single- and multi-point mutation subsets, achieving superior performance in 5 out of 7 metrics when evaluated across all mutations. (2) On this non-superfamily leakage test set, traditional force field simulators (e.g., the SoTA flex ddG) are surpassed by USP-ddG not only in computational efficiency but also in predictive accuracy. (3) Without utilizing any additional pre-training data, USP-ddG exceeds all pre-training based methods across all metrics, suggesting that our unified structural framework is intrinsically more effective for *ΔΔG* prediction than approaches leveraging external pre-training knowledge. (4) USP-ddG achieves clear improvements over BA-DDG, the strongest baseline, which has substantially outperformed other methods under PPI-disjoint cross-validation. (5) Finally, USP-ddG demonstrates the largest gains on multi-point mutations, a challenging due to the complexity of epistatic effects, thereby highlighting its strong practical applicability.

**Table 1:**
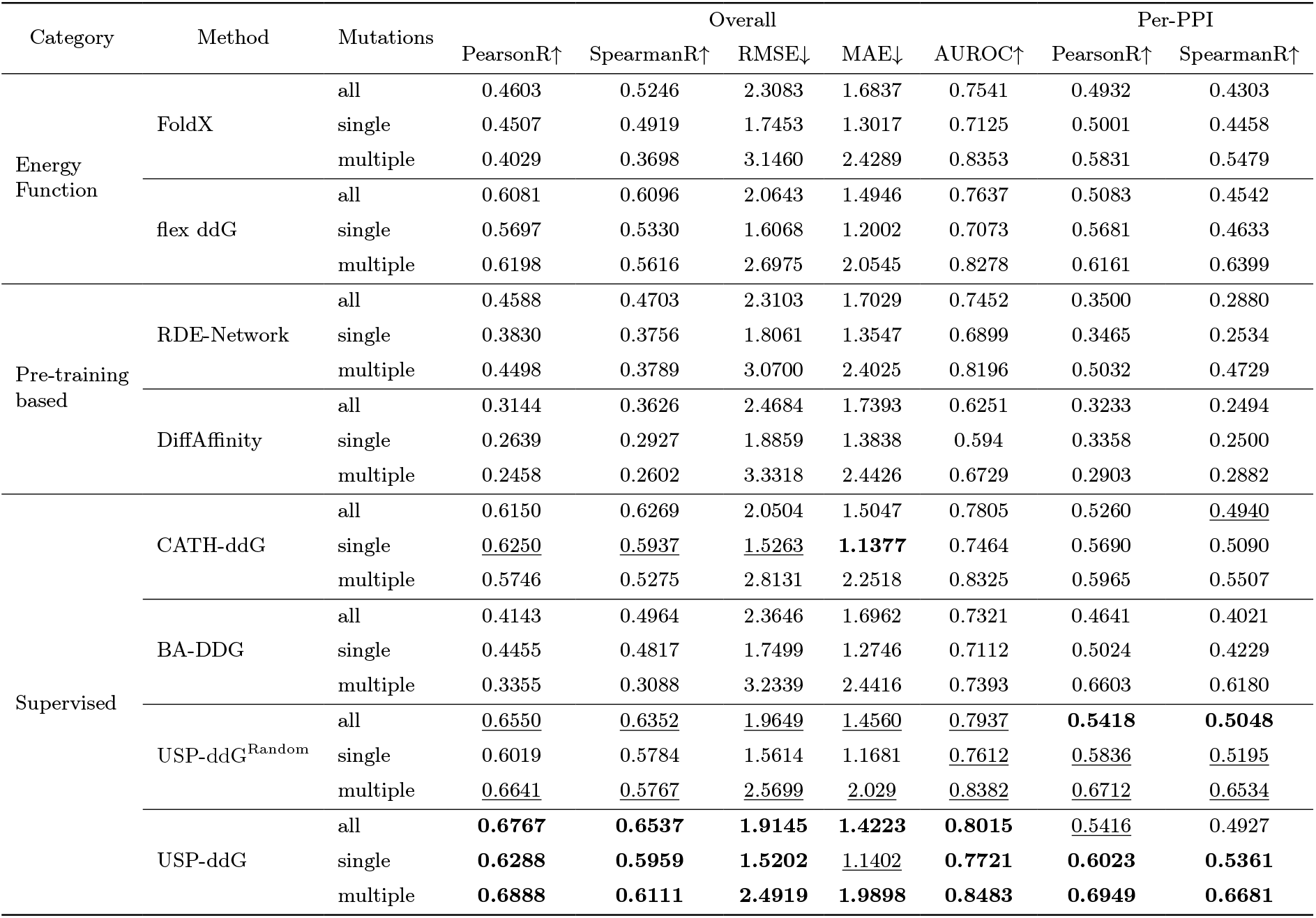
Performance on the hold-out CATH test set under all-, single-, multi-point mutations.

### 4.2 Evaluation on the antibody CR2621 against H1N1

CR6261 is a broadly neutralizing antibody (bnAb) against influenza A. Phillips et al. [31] systematically measured the *K*_*D*_ of combinatorially complete mutational libraries for 14 heavy-chain mutations, reconstructing all possible evolutionary intermediates to the germline sequence. Leveraging these experimental data, AbBiBench curated a benchmark of 1,887 mutants targeting A/New Caledonia/20/1999 (H1N1) for evaluating model performance in designing high-affinity antibodies (PDB ID: 3GBN) [47]. Crucially, none of these mutants appears in any public training databases, ensuring the absence of mutation-level data leakage and enabling reliable assessment of generalization capability. Accurately predicting *ΔΔG* for the variants of antibody CR6261 presents a challenge due to the prevalence of multi-point mutations and epistatic effects. Epistasis introduces non-additive interactions between mutations, which not only constrains the evolutionary paths accessible under selective pressure but also complicates the difficulty of accurately predicting *ΔΔG*. The results in Table 2 show that USP-ddG achieves the best performance measured by PearsonR and SpearmanR, with PearsonR and SpearmanR values exceeding the SoTA baseline BA-DDG by 12.86% and 10.78%, respectively, demonstrating its superior generalization ability in capturing epistatic effects.

**Table 2:** Evaluation of the binding affinity of CR6261 against H1N1.

| Category | Method | PearsonR↑ | SpearmanR↑ |
| --- | --- | --- | --- |
| Energy Function | FoldX | 0.577 | 0.593 |
|  | flex ddG | 0.515 | 0.516 |
| Pre-training based | RDE-Network | 0.370 | 0.422 |
|  | DiffAffinity | 0.351 | 0.323 |
| Supervised | CATH-ddG | 0.499 | 0.502 |
|  | BA-DDG | 0.583 | 0.566 |
|  | USP-ddG<sup>Random</sup> | 0.608 | 0.584 |
|  | USP-ddG | **0.658** | **0.627** |

### 4.3 Evaluation on the antibody trastuzumab against HER2

The HER2 binders test set, sourced from [6,36] (PDB ID: 1N8Z), comprises 419 variants featuring de novo designed complementarity determining region (CDR) loops. These variants were computationally designed to enhance the quality and controllability of the trastuzumab antibody using generative AI and have been experimentally validated by surface plasmon resonance (SPR), thus establishing the reliability for benchmarking AI-based antibody design methods.The antibody trastuzumab variants exhibit a high average edit distance of 7.6, presenting a significant challenge for *ΔΔG* predictors which are primarily trained on the low edit distance SKEMPI v2.0 dataset [6]. As shown in Table 3, USP-ddG achieves the best performance on this independent test set, with PearsonR and SpearmanR values that are 3.79% and 3.44% higher, respectively, than the SoTA baseline CATH-ddG. In contrast, traditional energy function methods such as flex ddG and FoldX are substantially outperformed by machine learning-based approaches on this HER2 binder test set.

**Table 3:** Evaluation on the binding affinity of HER2 binders.

| Category | Method | PearsonR↑ | SpearmanR↑ |
| --- | --- | --- | --- |
| Energy Function | FoldX | 0.246 | 0.329 |
|  | flex ddG | 0.301 | 0.339 |
| Pre-training based | RDE-Network | 0.465 | 0.481 |
|  | DiffAffinity | 0.590 | 0.524 |
|  | GearBind+Ensemble<sup>a</sup> | 0.579 | 0.608 |
| Supervised | CATH-ddG | 0.607 | 0.639 |
|  | BA-DDG | 0.326 | 0.362 |
|  | USP-ddG<sup>Random</sup> | 0.619 | 0.656 |
|  | USP-ddG | **0.630** | **0.661** |

<sup>a</sup> Results are from GearBind [6].

### 4.4 Ablation study

To elucidate the contribution of each component in USP-ddG, we performed an ablation study on the hold-out CATH test set. The model was systematically evaluated by progressively deactivating individual modules, followed by retraining and assessment. As illustrated in Table 4, we observed consistent declines in correlation metrics, underscoring the importance of each module in achieving optimal performance. Our results indicate that the inverse folding log *p*( 𝒮_𝒜ℬ_ | 𝒳^FX^) and CATH-ddG log *p*_*θ*_( 𝒮_𝒜ℬ_, 𝒳^RX^) modules contribute nearly equally to model performance. Specifically, CATH-ddG tends to offer a slightly greater contribution in scenarios involving multiple-point mutations, whereas inverse folding contributes more in single-point mutation cases. In contrast, the impact of data efficacy module (w/o Sampler) is less pronounced than that of the MoE module. Nevertheless, we observed that incorporating data efficacy improves generalization, which is evidenced in the case studies discussed above, thereby confirming its positive effect on model robustness.

**Table 4:**
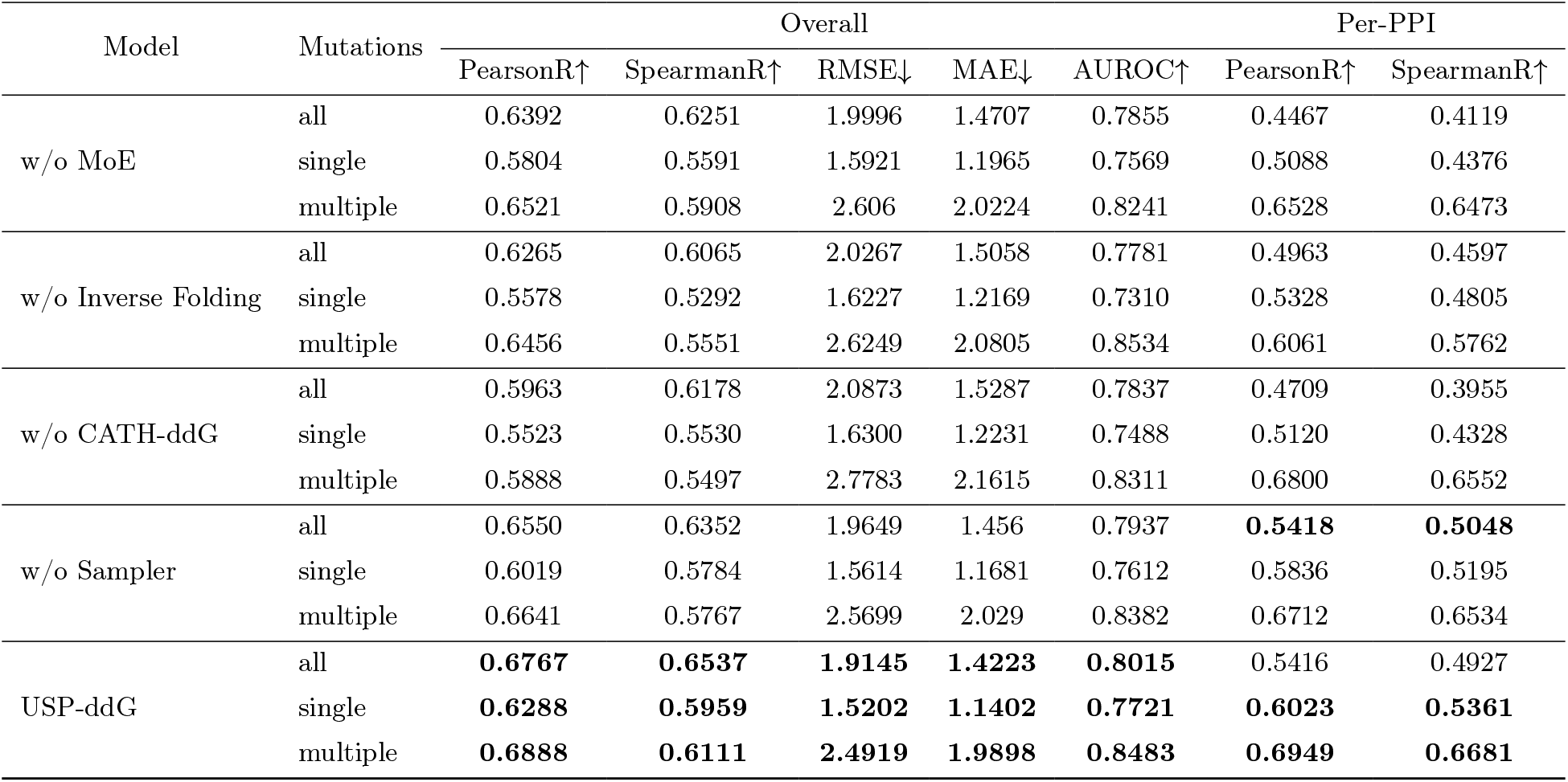
Impact of individual module exclusion on performance drop in the hold-out CATH test set.

### 4.5 Domain Specialization

Following [29], domain specialization characterizes the functional preference of expert ℰ_*i*_ for CATH domain 𝒞_*j*_, representing the percentage of tokens from 𝒞_*j*_ assigned to ℰ_*i*_, with 100% indicating exclusive dedication and 0% identifying prunable experts for that domain. Fig. 2 reveals that many experts are significantly activated deviating from the uniform routing baseline on specific CATH domains. A notable example is the EphA2-SAM/SHIP2-SAM complex, where C-terminal sterile alpha motifs (SAM) form a heterodimer that negatively regulates receptor endocytosis and degradation [41], resulting in the 2nd and 3rd experts in each layer specialize over 50%. This reflects that the domain specialization in USP-ddG results in low functional overlap, with dedicated experts exhibiting distinct domain preferences, thereby effectively alleviating structural overlaps among protein domains as intended. In addition, as the E9 DNase/Im9 and E9 DNase/Im2 complexes share similar CATH domains mediated by structurally analogous PPIs, they exhibit closely aligned domain specializations. Likewise, EphA3/ephrinA5 and EphA4/ephrinB2 PPIs also exhibit similar domain specializations. Consequently, as shown in Fig. 2 below, the MoE framework demonstrates the ability to precisely delineate these domain-specific features, this highlights its advantage in identifying determinants of PPIs and showcasing its potential to advance the development of protein domain representation learning.

**Fig. 2:**
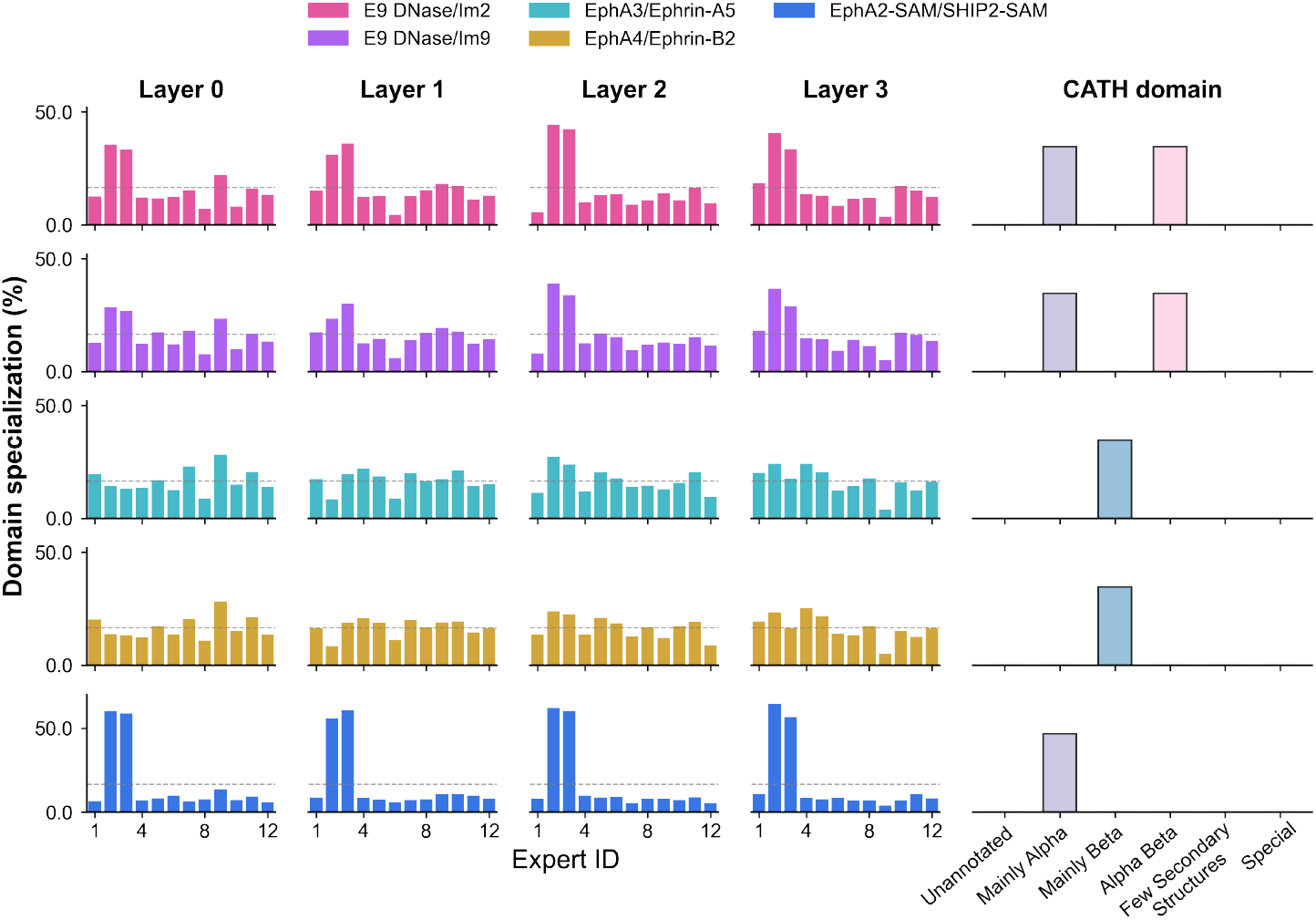
Domain specialization of USP-ddG. We visualize the routing distribution of tokens from different protein domains across each layer to the 12 experts for the five PPIs in the hold-CATH test set, considering the top-2 active experts per token. The horizontal gray dashed lines represent the uniform routing baseline of 16.67% (2*/*12) per expert, which results from activating 2 out of 12 experts each token in every layer, providing reference for assessing the learned domain specialization.

## 5 Discussion

Predicting *ΔΔG* for PPIs with the unified structural framework is not only a critical computational task, but also facilitates to a comprehensive understanding of proteins as the ultimate executors of cellular structures and functions. However, current methods lack a systematic consensus on mutation-induced conformational changes and often overlook the necessity of establishing a unified structural paradigm. In this work, we present USP-ddG, a pioneering dual-channel architecture that significantly advances the prediction of the changes in binding free energy. By integrating structure-fixed and -relaxed assumptions via a hybrid layer of FFN and MoE, coupled with data efficacy training strategy, USP-ddG outperforms existing approaches and offers a robust tool for *ΔΔG* prediction.

Extensive experimental results demonstrate that USP-ddG achieves outstanding performance across various metrics. Our ablation studies further confirm the importance of each component to the overall performance. While USP-ddG represents a significant advancement, certain limitations remain. Two principal bottlenecks currently limit the accuracy and scalability of *ΔΔG* prediction. First, the scarcity of high-resolution complex structures necessitates the use of structure prediction tools such as AlphaFold3 [1]. Second, the reliance on tools like FoldX for generating mutant variants incurs computationally burden when applied at scale, emphasizing the need for end-to-end approaches capable of inferring mutational effects directly from wild-type structures. Addressing these challenges would enhance the efficiency and scalability of the proposed *in silico* antibody affinity maturation pipeline, thereby advancing therapeutic antibody engineering and contributing to disease therapy and biomedical research.

