## Appendix for "USP-ddG: A Unified Structural Paradigm with Data Efficacy and Mixture-of-Experts for Predicting Mutational Effects on Protein-Protein Interactions"

In Appendix A, we describe how to use inverse folding model to predict $\Delta\Delta G$. In Appendix B, we provide a detailed derivation of the binding free energy $\Delta G^{FX}$ under the structure-fixed hypothesis. In Appendix C and Appendix D, we provide a detailed description of the seven metrics used to evaluate $\Delta\Delta G$ prediction performance and the parameter settings in our proposed USP-ddG model. In Appendix E, we discuss the experiments conducted on the SARS-CoV-2 RBD to hACE2.

### A $\Delta\Delta G$ Prediction Using Inverse Folding

From the definition of $\Delta G$, previous work such as BA-DDG [2] interprets $G_{\text{bnd}}$ and $G_{\text{unbnd}}$ as the conditional probabilities of the complex structure being in the bound conformation $\mathcal{X}_{\text{bnd}}$ or unbound conformation $\mathcal{X}_{\text{unbnd}}$, respectively, given its sequence $\mathcal{S}_{\mathcal{AB}}$. Thus, $\Delta G$ is defined by the Boltzmann distribution [1]:

$$
\begin{aligned}
G_{\text{bnd}} &= -k_B T \cdot \log p(\mathcal{X}_{\text{bnd}} \mid \mathcal{S}_{\mathcal{AB}}), && (1)\\
G_{\text{unbnd}} &= -k_B T \cdot \log p(\mathcal{X}_{\text{unbnd}} \mid \mathcal{S}_{\mathcal{AB}}), && (2)\\
\Delta G &= G_{\text{bnd}} - G_{\text{unbnd}}, && (3)\\
&= -k_B T \cdot \left(\log p(\mathcal{X}_{\text{bnd}} \mid \mathcal{S}_{\mathcal{AB}}) - \log p(\mathcal{X}_{\text{unbnd}} \mid \mathcal{S}_{\mathcal{AB}})\right), && (4)
\end{aligned}
$$

where $k_B$ is the Boltzmann constant and $T$ is the thermodynamic temperature. Subsequently, applying Bayes' theorem and assuming the protein backbone structure remains unchanged before and after mutation (i.e. $\mathcal{X}^{\text{mut}} = \mathcal{X}^{\text{wt}}$), $\Delta\Delta G$ can be simplified via the thermodynamic cycle to:

$$
\Delta\Delta G = -k_B T \cdot \left(\log \frac{p(\mathcal{S}^{\text{mut}}_{\mathcal{AB}} \mid \mathcal{X}^{\text{mut}}_{\text{bnd}})}{p(\mathcal{S}^{\text{mut}}_{\mathcal{AB}} \mid \mathcal{X}^{\text{mut}}_{\text{unbnd}})} - \log \frac{p(\mathcal{S}^{\text{wt}}_{\mathcal{AB}} \mid \mathcal{X}^{\text{wt}}_{\text{bnd}})}{p(\mathcal{S}^{\text{wt}}_{\mathcal{AB}} \mid \mathcal{X}^{\text{wt}}_{\text{unbnd}})}\right), \quad (5)
$$

where the conditional probability of the unbound state, $p(\mathcal{S}^{\text{mut}}_{\mathcal{AB}} \mid \mathcal{X}^{\text{mut}}_{\text{unbnd}})$, is approximated as the product of the conditional probabilities for the individual monomers, that is $p(\mathcal{S}^{\text{mut}}_{\mathcal{AB}} \mid \mathcal{X}^{\text{mut}}_{\text{unbnd}}) \approx p(\mathcal{S}^{\text{mut}}_{\mathcal{A}} \mid \mathcal{X}^{\text{mut}}_{\mathcal{A}}) \cdot p(\mathcal{S}^{\text{mut}}_{\mathcal{B}} \mid \mathcal{X}^{\text{mut}}_{\mathcal{B}})$. Inverse folding models, such as ProteinMPNN, are trained to estimate the negative log-likelihood scores of a protein sequence $\mathcal{S}$ given its backbone structure $\mathcal{X}$, thereby providing an approximation of $p(\mathcal{S} \mid \mathcal{X})$. Consequently, all conditional probabilities in the Equation 5 above can be directly derived from the negative log-likelihood scores generated by ProteinMPNN.

### B Structure-fixed Assumption ($\Delta G^{\text{FX}}$)

Under the structure-fixed hypothesis, the binding free energy $\Delta G^{\text{FX}}$ can be derived from the Boltzmann distribution [1], and expressed in terms of the probabilities of the bound state $p^{\text{FX}}_{\text{bnd}}$ and the unbound state $p^{\text{FX}}_{\text{unbnd}}$:

$$
\begin{aligned}
\Delta G^{\text{FX}} &= G^{\text{FX}}_{\text{bnd}} - G^{\text{FX}}_{\text{unbnd}}, && (6)\\
&= -k_B T \cdot \left(\log p^{\text{FX}}_{\text{bnd}} - \log p^{\text{FX}}_{\text{unbnd}}\right), && (7)\\
&= -k_B T \cdot \left(\log p(\mathcal{S}_{\mathcal{AB}}, \mathcal{X}^{\text{FX}}_{\text{bnd}}) - \log p(\mathcal{S}_{\mathcal{AB}}, \mathcal{X}^{\text{FX}}_{\text{unbnd}})\right). && (8)
\end{aligned}
$$

Applying Bayes' theorem, similar to the EBM-DDG approach [5], this simplifies to:

$$\begin{aligned}\Delta G^{\mathrm{FX}} &= -k_B T \cdot \left(\log p(\mathcal{S}_{\mathcal{AB}}, \mathcal{X}^{\mathrm{FX}}_{\mathrm{bnd}}) - \log p(\mathcal{S}_{\mathcal{AB}}, \mathcal{X}^{\mathrm{FX}}_{\mathrm{unbnd}})\right), &(9)\\ &= -k_B T \cdot \left(\log p(\mathcal{S}_{\mathcal{AB}} \mid \mathcal{X}^{\mathrm{FX}}_{\mathrm{bnd}}) p(\mathcal{X}^{\mathrm{FX}}_{\mathrm{bnd}}) \right.\\ &\qquad \left. - \log p(\mathcal{S}_{\mathcal{AB}} \mid \mathcal{X}^{\mathrm{FX}}_{\mathrm{unbnd}}) p(\mathcal{X}^{\mathrm{FX}}_{\mathrm{unbnd}})\right), &(10)\\ &= -k_B T \cdot \left(\log \frac{p(\mathcal{S}_{\mathcal{AB}} \mid \mathcal{X}^{\mathrm{FX}}_{\mathrm{bnd}})}{p(\mathcal{S}_{\mathcal{AB}} \mid \mathcal{X}^{\mathrm{FX}}_{\mathrm{unbnd}})} + \log \frac{p(\mathcal{X}^{\mathrm{FX}}_{\mathrm{bnd}})}{p(\mathcal{X}^{\mathrm{FX}}_{\mathrm{unbnd}})}\right). &(11)\end{aligned}$$

The unbound state probability is approximated as $p(\mathcal{S}_{\mathcal{AB}} \mid \mathcal{X}^{\mathrm{FX}}_{\mathrm{unbnd}}) \approx p(\mathcal{S}_{\mathcal{A}} \mid \mathcal{X}^{\mathrm{FX}}_{\mathcal{A}}) \cdot p(\mathcal{S}_{\mathcal{B}} \mid \mathcal{X}^{\mathrm{FX}}_{\mathcal{B}})$. Furthermore, leveraging the common assumption in protein sequence design that the sequence is largely determined by the backbone structure, we approximate:

$$\begin{aligned}\log p(\mathcal{S}_{\mathcal{AB}} \mid \mathcal{X}^{\mathrm{FX}}_{\mathrm{bnd}}) &= \log p(\mathcal{S}_{\mathcal{AB}} \mid \{\mathcal{X}^{\mathrm{FX}}_{\mathrm{bnd,BB}}, \mathcal{X}^{\mathrm{FX}}_{\mathrm{bnd,SC}}\}) \approx \log p(\mathcal{S}_{\mathcal{AB}} \mid \mathcal{X}^{\mathrm{FX}}_{\mathrm{bnd,BB}}), &(12)\\ \log p(\mathcal{S}_{\mathcal{A}} \mid \mathcal{X}^{\mathrm{FX}}_{\mathcal{A}}) &= \log p(\mathcal{S}_{\mathcal{A}} \mid \{\mathcal{X}^{\mathrm{FX}}_{\mathcal{A},\mathrm{BB}}, \mathcal{X}^{\mathrm{FX}}_{\mathcal{A},\mathrm{SC}}\}) \approx \log p(\mathcal{S}_{\mathcal{A}} \mid \mathcal{X}^{\mathrm{FX}}_{\mathcal{A},\mathrm{BB}}), &(13)\\ \log p(\mathcal{S}_{\mathcal{B}} \mid \mathcal{X}^{\mathrm{FX}}_{\mathcal{B}}) &= \log p(\mathcal{S}_{\mathcal{B}} \mid \{\mathcal{X}^{\mathrm{FX}}_{\mathcal{B},\mathrm{BB}}, \mathcal{X}^{\mathrm{FX}}_{\mathcal{B},\mathrm{SC}}\}) \approx \log p(\mathcal{S}_{\mathcal{B}} \mid \mathcal{X}^{\mathrm{FX}}_{\mathcal{B},\mathrm{BB}}). &(14)\end{aligned}$$

Thus, the conditional probabilities $p(\mathcal{S}_{\mathcal{AB}} \mid \mathcal{X}^{\mathrm{FX}}_{\mathrm{bnd}})$, $p(\mathcal{S}_{\mathcal{A}} \mid \mathcal{X}^{\mathrm{FX}}_{\mathcal{A}})$, and $p(\mathcal{S}_{\mathcal{B}} \mid \mathcal{X}^{\mathrm{FX}}_{\mathcal{B}})$ are provided by a pre-trained inverse folding model, denoted as $\Delta G_{\mathrm{ProteinMPNN}}$.

And the second term in Equation 11, $\log \frac{p(\mathcal{X}^{\mathrm{FX}}_{\mathrm{bnd}})}{p(\mathcal{X}^{\mathrm{FX}}_{\mathrm{unbnd}})}$, which corresponds to the binding free energy between the bound and unbound states of the full complex, is estimated by EBM-DDG under the structure-fixed and backbone-only assumption using the diffusion-based score-matching model DSMBind [3]. In contrast, by rethinking the thermodynamic cycle definition of $\Delta\Delta G$ and incorporating the consideration of protein side-chain structures, we propose to approximate this energy term under the empirical force field FoldX. This is achieved using its "Interaction Energy" term, denoted as $\Delta G_{\mathrm{FoldX}}$:

$$\begin{aligned}\Delta G_{\mathrm{FoldX}} &= -k_B T \cdot \log \frac{p(\mathcal{X}^{\mathrm{FX}}_{\mathrm{bnd}})}{p(\mathcal{X}^{\mathrm{FX}}_{\mathrm{unbnd}})}, &(15)\\ &= -k_B T \cdot \log \frac{p(\{\mathcal{X}^{\mathrm{FX}}_{\mathrm{bnd,BB}}, \mathcal{X}^{\mathrm{FX}}_{\mathrm{bnd,SC}}\})}{p(\{\mathcal{X}^{\mathrm{FX}}_{\mathrm{unbnd,BB}}, \mathcal{X}^{\mathrm{FX}}_{\mathrm{unbnd,SC}}\})}. &(16)\end{aligned}$$

### C Evaluation Metrics

We employ a comprehensive set of seven metrics to evaluate $\Delta\Delta G$ prediction performance. Five overall metrics: (1) Pearson correlation coefficient; (2) Spearman's rank correlation coefficient; (3) Root Mean Squared Error (RMSE); (4) Mean Absolute Error (MAE); and (5) AUROC, where mutations are classified based on the sign of ground-truth $\Delta\Delta G$ values. Given the practical importance of correlation within specific PPIs, we report two additional PPI-wise metrics. Mutations are grouped by PPI, and the average Pearson and Spearman correlation coefficients across PPIs are reported as Per-PPI correlation metrics.

### D Parameter Settings

We implemented USP-ddG with PyTorch and optimized the model parameters using the Adam algorithm with the default parameters ($\beta_1 = 0.9, \beta_2 = 0.999$), combined with cyclical cosine annealing scheduler for adaptive learning rate adjustment. We configured the MoE layer with 12 experts, activating the top-2 for each input token, along with the LoRA MoE-Adapter of rank $r = 64$ and model dimension $d_{\mathrm{model}} = 128$. The batch size is set to 12 and all experiments are performed on an NVIDIA GeForce RTX 2080 Ti GPU. Following the CATH-ddG protocol, we employed a data augmentation technique that incorporates hybrid Gaussian noise for the structure-relaxed channel, as shown in Fig. 1 (a). Specifically, we applied distinct noise levels to the atomic coordinates, with std = 0.1 Å for the backbone atoms and std = 0.2 Å for the side chain atoms. In addition, we crop each protein structure into patches of $L_{\mathrm{patch}} = 256$ residues by selecting mutation sites as the anchor and subsequently choosing its 255 nearest neighbors according to $C_\beta$-$C_\beta$ distances.

### E Evaluation on the SARS-CoV-2 RBD to hACE2

The evolution of the SARS-CoV-2 spike receptor-binding domain (RBD) is shaped by epistasis, leading to accumulated substitutions that enhance ACE2 binding affinity and enable escape from antibody recognition. To comprehensively quantify how epistasis alters mutational effects, [6] performed deep mutational scans (DMS) on the ancestral Wuhan-Hu-1 RBDs of all single-point mutations. Following [4], we benchmark predictions against 285 single-point mutations across 15 critical RBD sites. As shown in Table 1, USP-ddG achieves a PearsonR of 0.598 between the experimental and predicted $\Delta\Delta G$, markedly surpassing all other baseline methods.

Table 1: Evaluation of the binding affinity between SARS-CoV-2 RBD and hACE2.

| Method | FoldX[a] | RDE-Network[a] | CATH-ddG[a] | BA-DDG[a] | USP-ddG$^{\mathrm{Random}}$ | USP-ddG |
|---|---|---|---|---|---|---|
| PearsonR↑[b] | 0.385 | 0.403 | 0.579 | 0.473 | 0.572 | **0.598** |

[a] Results are from released tools or source code.
[b] The **bold** value indicates the best result.
